# Refining the Bio-manufacturing of Microalgae-derived Extracellular Vesicles as a Potential Nanotherapeutic for Osteoarthritis

**DOI:** 10.1101/2025.10.30.685512

**Authors:** Meng Wang, Sara Gil Izquierdo, Aylin Kara Özenler, Jaqueline Lourdes Rios, Mylène de Ruijter, Debby Gawlitta, Jos Malda, Kenny Man

## Abstract

Osteoarthritis (OA) is a degenerative joint disease marked by oxidative stress, chronic inflammation, and cartilage degradation. Current treatments are limited and fail to address the underlying disease mechanisms. Extracellular vesicles (EVs) have emerged as promising nanotherapeutics; however, mammalian-derived EVs face cost and scalability challenges. Microalgae represent a sustainable alternative, yet their potential as EV biofactories for regenerative medicine remains unknown. This study aims to refine the bio-manufacturing of microalgae EVs as a next-generation nanotherapeutic for OA.

Microalgae species (*Chlorella sorokiniana, Synechococcus* sp.*, Leptolyngbya* sp.*, Chlamydomonas reinhardtii CC1690*) were screened under varying photoperiods (0h, 16h, 24h light/day) to assess the influence on viability, growth and EV production. EVs were characterized by transmission electron microscopy, nanoparticle tracking analysis, protein quantification and immunoblotting. Their antioxidant capacity, cellular recruitment, and therapeutic efficacy were evaluated in a cytokine-induced OA-like *in vitro* model.

Our findings demonstrated that microalgae growth and EV yield were highly light-dependent, with all species maintaining high viability (>80%) across different photoperiods. Notably, *Leptolyngbya* sp. (*Leptolyngbya*) exhibited the fastest growth and highest EV yield under extended illumination, producing EVs with strong antioxidant activity. *Leptolyngbya-*derived EVs (Lepto-EVs) enhanced the proliferation and migration of human bone marrow-derived mesenchymal stromal cells and provided protection against matrix degradation within a cytokine-induced OA-like model.

These findings position microalgae, particularly *Leptolyngbya*, as a highly scalable and sustainable producer of therapeutic EVs. Lepto-EVs offer a potent, cell-free nanotherapeutic exhibiting anti-catabolic properties alleviating cytokine-induced matrix degradation in an OA-like milieu, establishing microalgae EVs as a promising, cost-effective frontier in regenerative nanomedicine.

**Graphical abstract:** 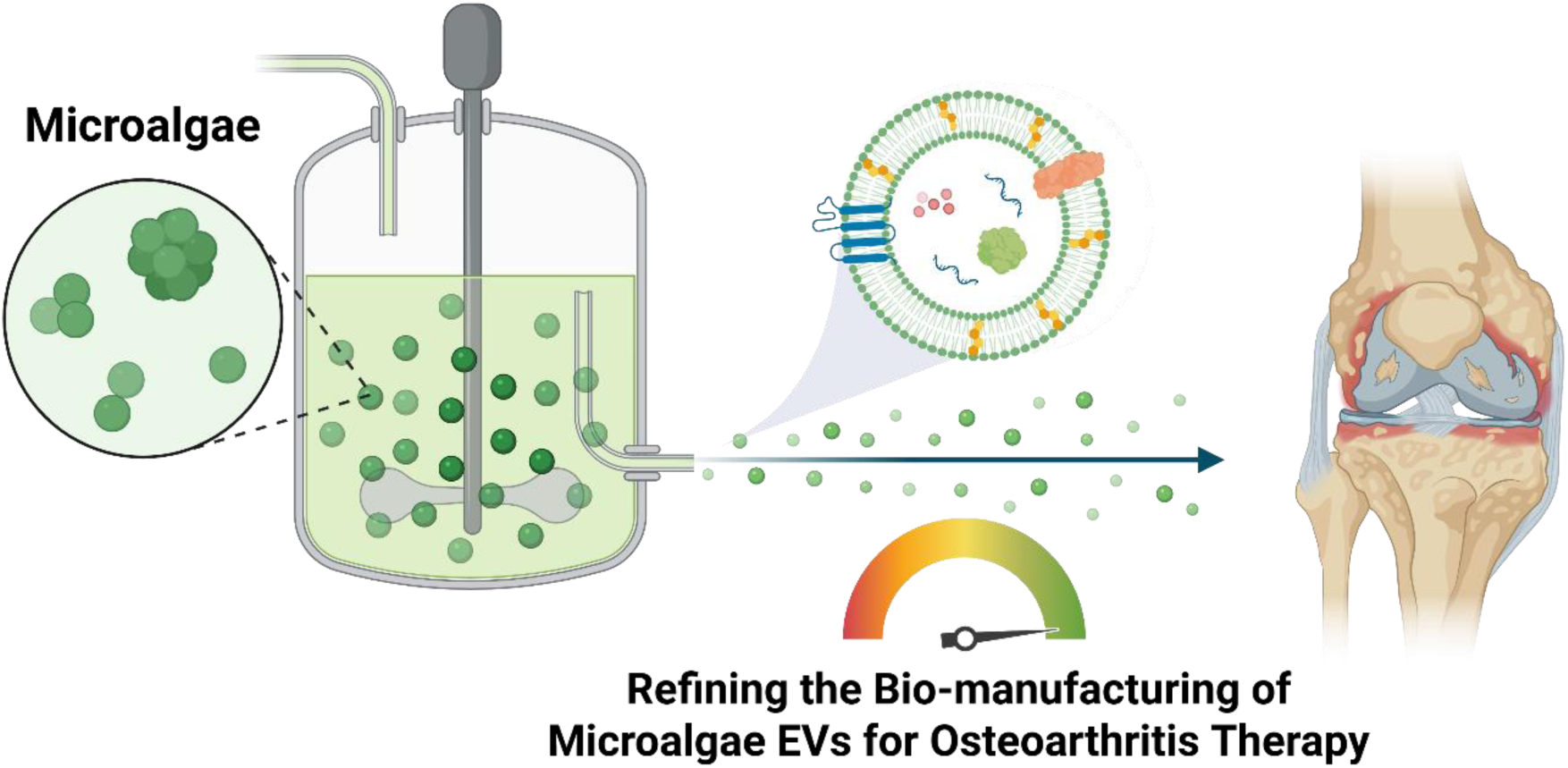

## 1. Introduction

Osteoarthritis (OA) is a chronic degenerative joint disease, affecting millions and causing significant disability. Characterized by pain, dysfunction, and joint deformity, OA places a significant burden on healthcare systems and severely diminishes patients’ quality of life ^1,2^. The global prevalence of OA was estimated at 7.96% of the global population in 2020 ^3^, which is projected to rise due to our growing, ageing population, thus incurring substantial economic costs related to treatment, rehabilitation, and lost productivity ^4,5^. OA is a multifactorial disorder shaped by a complex interplay of intrinsic and extrinsic factors, including age, sex, genetic predisposition, obesity, prior joint injury, diet, and lifestyle ^6^. There is currently no standard-of-care for OA, where disease management relies on conservative approaches such as corticosteroid injections, non-steroidal anti-inflammatory drugs, physical therapy, or lifestyle changes ^7^. Surgical treatments such as microfracture and mosaicplasty can alleviate pain ^8^, however often result in unsatisfactory long-term outcomes ^1,9^. Currently there is no cure for OA, emphasizing the need for novel therapies that target the underlying cause of the disease. While the precise etiology of OA remains elusive, accumulating evidence points to a confluence of factors, including chronic inflammatory signaling cascades ^10^, oxidative stress ^11^, chondrocyte apoptosis ^12^, and dysregulated energy metabolism ^13^. Thus, the development of novel therapeutic interventions targeting the modulation of these factors represents a promising avenue for the development of disease-modifying OA treatments.

Cell-based strategies utilizing mesenchymal stem/stromal cells (MSCs) have shown promise due to their immunomodulatory capabilities ^14,15^. Although promising, their clinical translation is hampered by significant hurdles, including poor cell survival, immune rejection, ethical issues, complex regulatory approval, high production cost and potential tumour formation ^16,17^. Nanotechnology offers a powerful solution to these challenges, enabling targeted delivery, improved biocompatibility, the ability to traverse biological barriers (*e.g*., blood-brain barrier), off-the-shelf storage, and minimized adverse immune responses ^18,19^. Extracellular vesicles (EVs) are emerging as a promising nanoscale therapy for musculoskeletal repair, overcoming limitations associated with cell-based therapies. EVs are cell-secreted lipid nanoparticles, which contain a package of bioactive molecules (*i.e.*, proteins, metabolites, nucleic acids) and play a crucial role in intercellular communication ^19–21^. By eliminating the risks of uncontrolled differentiation, tumorigenicity, and poor survival associated with cell therapies, EV-based therapies offer an exciting prospect for a readily available, off-the-shelf OA treatment ^22–24^. Moreover, EVs can readily encapsulate and deliver various therapeutic molecules (*e.g.,* anti-inflammatory drugs, growth factors) directly to the defect site, enhancing their efficacy and reducing side effects ^25,26^. Moreover, the economic advantage of EV-based approaches in comparison to MSC therapies has been reported, with a dose of MSC EVs (10^11^ particles) costing up to €3,082, whilst clinical doses of MSCs, can cost up to €42,673 depending on dose size and production scale ^27^.

Despite their therapeutic and economic appeal, the clinical translation of mammalian cell-derived EVs remains limited by several practical challenges, including donor variability, low production yields, labour-intensive isolation procedures, and difficulties in large-scale manufacturing and long-term storage ^17,28^. In contrast, microalgae constitute a sustainable and renewable source of bioactive compounds widely utilized in the health, cosmetics, and food industries ^29,30^. Microalgae are photosynthetic autotrophic microorganisms, rich in bioactive metabolites, such as pigments, vitamins, antioxidants ^31,32^. Critically, algae cultivation is scalable, cost-effective, and environmentally friendly, overcoming the limitations of mammalian cell-based production ^33,34^. Therefore, microalgae could provide a robust renewable platform for generating therapeutic EVs at clinically relevant scales. Despite emerging interest in the biomedical applications of microalgae and their derivatives, their potential as a platform for producing therapeutic EVs remains underexplored. In particular, the refinement of algal cultivation strategies to optimise EV yield and quality has received little attention. Given that microalgae are highly sensitive to environmental conditions, factors such as light intensity, nutrient availability, and temperature can significantly influence vesicle production ^35,36^, critical variables that must be controlled to meet clinical-grade standards.

This study aims to optimize the biomanufacturing of microalgal EVs and to evaluate these as a potential next generation nanotherapeutics for OA. The research specifically focuses on selecting a potent microalgal species with a high EV production yield, evaluating the effects of different light regimes on algal growth and EV biogenesis, and determining how medium harvest frequency influences EV yield and quality (Fig. 1). The most productive and scalable species was subsequently chosen for further investigation, where the potency of its EVs was assessed for their ability to mitigate inflammation-induced cartilage-like ECM degradation in a cytokine-induced OA-like *in vitro* model.

**Figure 1.**
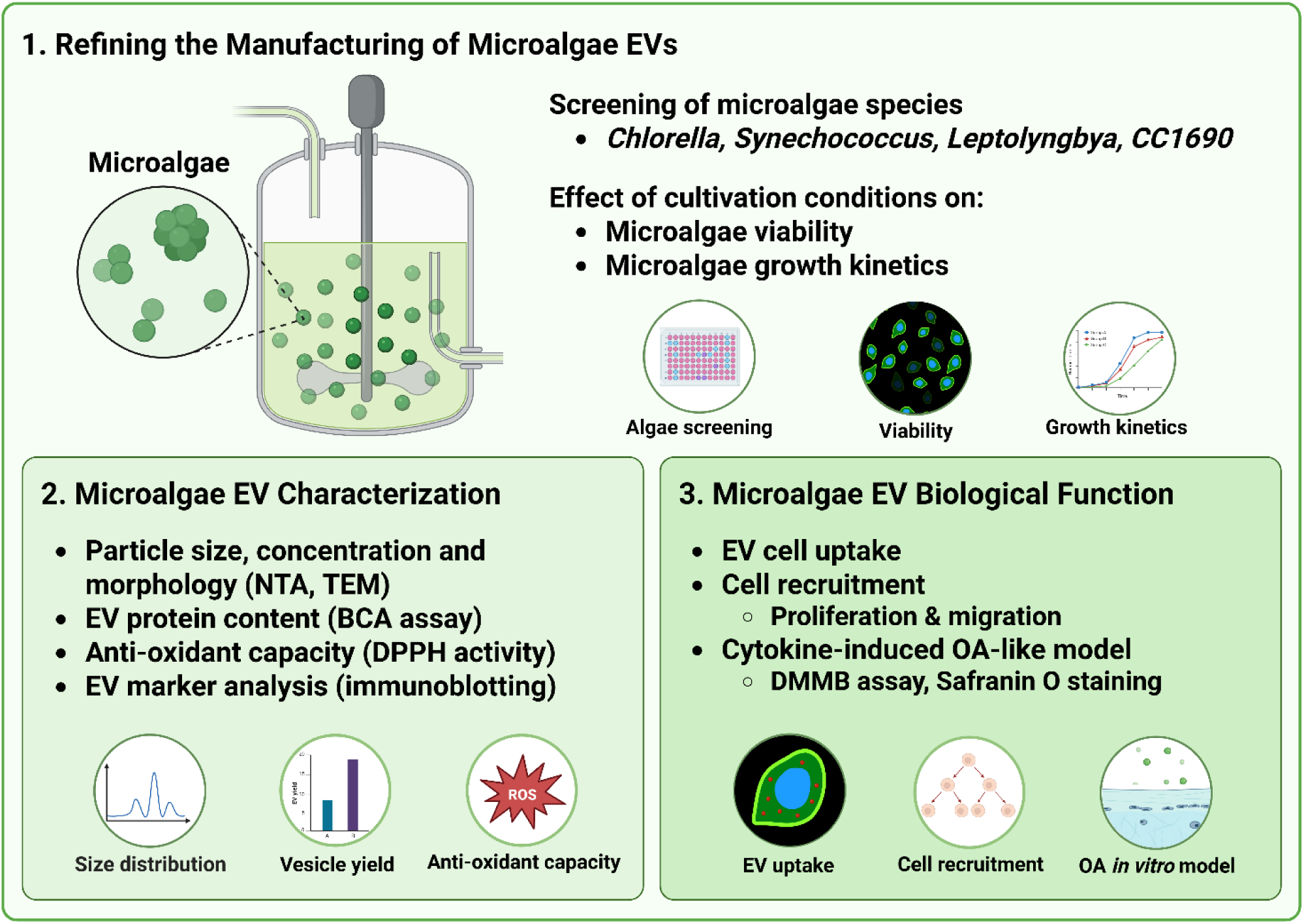
Experimental overview investigating the effect of biomanufacturing conditions on microalgae EV production and therapeutic potency. 1) The influence of different manufacturing conditions (i.e., light exposure, harvesting frequency) on microalgae viability, growth kinetics, and yield. *Chlorella sorokiniana* (*Chlorella*)*, Synechococcus* sp. (*Synechococcus*)*, Leptolyngbya* sp. (*Leptolyngbya*)*, Chlamydomonas reinhardtii CC1690* (*CC1690*). 2) EVs were isolated from microalgae over a 2-week period, and the nanoparticles were characterized by their size distribution, morphology, protein content, antioxidant capacity, and EV marker expression. 3) Investigating the influence of microalgae EV treatment on recipient cell uptake, proliferation, migration, and anti-catabolic properties in an OA-like *in vitro* model.

## 2. Materials and Methods

### 2.1. Microalgae cultivation

*Chlorella sorokiniana*, *Synechococcus* sp., *Leptolyngbya* sp., and *Chlamydomonas reinhardtii CC1690* with initial concentration of 25 × 10⁶ cells/mL were cultured separately in modified M8 medium, sterile artificial seawater medium, and modified Tris-Acetate-Phosphate (TAP) medium (Supplementary Table 1) on shakers (Multitron, Infors HT, Switzerland) with 150 rpm of agitation speed. All the microalgal cultivation media were supplemented with HEPES (20 mM) to stabilize pH. Bright white LED (LedstripKoning, HWCS600-03M) was used as artificial light source, and different photoperiods (0, 16 and 24h of light/day) were applied accordingly to stimulate microalgae growth. The light intensity was approximately 100 μmol photons‧m⁻²‧s⁻¹. All the microalgal cultures were maintained for 2 weeks, with medium changes occurring once or twice a week, unless otherwise stated. Briefly, microalgal cultures were harvested and centrifuged at 1200 g for 5 min, the resulting microalgae pellets were resuspended in fresh media and returned to the original flasks/shakers for further cultivation.

### 2.2 Microalgae growth kinetics and viability

To determine microalgal number and biomass, the optical density at 750 nm (OD₇₅₀) was measured (Supplementary Table 2, and 3). For each measurement, a total of 100 µL of diluted sample was placed into a 96 well microplate and read using a microplate reader (CLARIOstar Plus, BMG LABTECH, Ortenberg, Germany). To assess the growth kinetics of each microalgal species, normalized microalgal number (Nt) and biomass (Bt) were calculated via equation [1] and [2],

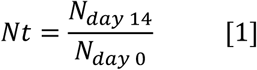

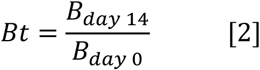

where N_t_ represents the normalized microalgal cell number, and N_day 14_ and N_day 0_ are the microalgal cell number after 2 weeks and the initial culture microalgal cell number, respectively. B_t_, represents the normalized microalgal dry biomass, and B_day 14_ and B_day 0_ are the microalgal biomass after 2 weeks and the biomass of initial microalgal used for cultivation, respectively.

The viable microorganisms can be traced by the autofluorescence of chlorophyll under red wavelength, which was detected with an excitation/emission of 640/700 nm. The non-viable microalgae were stained by SYTOX™ Orange (1 µM, Invitrogen™) and detected with an excitation/emission of 543/570 nm. At least 6 images of each cultivation condition were taken by Thunder microscopy (Leica Microsystems, Germany). Total area of both live and dead microalgae signals was quantified via Fiji ImageJ (https://imagej.net/software/fiji/) processing. Viability of microalgae was calculated as a percentage via equation [3], where TA_l_ and TA_d_ represent total area of live and dead microalgae separately.

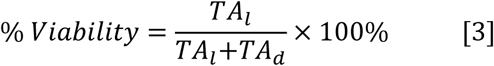

### 2.3 Microalgae EV isolation and characterization

The collected microalgae conditioned medium was centrifuged at 3000 g for 20 min to remove residual microalgal cellular debris. The supernatant was then filtered through a 0.22 µm membrane filter and stored at −80°C for subsequent EV isolation. The microalgae conditioned media was centrifuged at 10,000 g for 30 min at 4°C using an Optima XE-90 ultracentrifuge with a SW 32 Ti rotor (Beckman Coulter, USA). The resulting EV pellet was washed in sterile phosphate-buffered saline (PBS) and centrifuged following the same conditions. The EV pellet was then re-suspended in PBS and stored at −80°C until required.

To estimate EV particle size and concentration, five videos (30 sec each) were recorded using a NanoSight NS500 instrument (Malvern Panalytical, Great Malvern, UK) equipped with a sCMOS camera and a 405 nm blue laser, at 25°C. Samples were diluted between 1:100 and 1:1000 in PBS to obtain a particle number in the chamber between 20-100 to ensure optimal detection and tracking. Data was processed using NTA software version 3.4.

EVs were imaged with a FEI Tecnai 12 transmission electron microscope (Thermo Fisher Scientific, Waltham, MA, USA), operated at 80 kV high tension with image acquisition via the Tecnai User Interface software. The samples were placed in a carbon-coated copper grid and negatively stained. Briefly, grids were incubated in 2% uranyloxalate-acetate (pH 7) for 4 min and then in a solution containing 0.1% (w/v) methyl cellulose and 2% (w/v) uranyl acetate (pH 4) on ice for 5 min. After staining, excess solution was gently removed with filter paper and grids were air-dried before imaging.

Total EV protein concentration was determined using the Micro BCA Protein Assay Kit (Thermo Scientific, 23235) according to the manufacturer’s instructions. Briefly, 150 µL of sample and working reagent was added per well and incubated at 37°C for 2h. Absorbance was measured at 562 nm on a microplate reader (CLARIOstar Plus, BMG LABTECH, Ortenberg, Germany).

Immunoblotting analysis was conducted to assess the presence of EV-associated protein markers. The Exo-Check™ Antibody Array (System Biosciences, EXORAY200B-4), which detects EV markers (CD63, CD81, ALIX, FLOT1, ICAM1, EpCam, ANXA5, and TSG101), was employed following the manufactureŕs instructions. Briefly, 90 µg of EV protein was incubated within the membrane at 4°C overnight. Following which, the membrane was developed using the SuperSignal™ West Pico PLUS Chemiluminescent Substrate (Thermo Scientific, 34577) and images acquired with the ChemiDoc Imaging System (Bio-Rad Laboratories, Hercules, USA).

2,2-diphenyl-1-picrylhydrazyl (DPPH) assay was used to determine the total antioxidant potential of EV samples ^37,38^. 75 µL of DPPH• solution (50 µM in methanol) was added to 25 µL of EV samples containing 90 µg/mL of protein. Methanol was used for the baseline correction. 100 µg/mL of ascorbic acid was used as a positive control. The mixture was allowed to stand for 2 mins at room temperature. The change in absorbance was measured at 515 nm using the microplate reader (CLARIOstar Plus, BMG LABTECH, Ortenberg, Germany). The ability of DPPH• radical scavenging activity was calculated by using the following equation [4], where A0 is the absorbance of the control, and A1 is the absorbance of the EV sample.

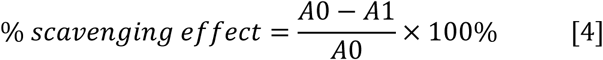

### 2.4. Mammalian cell cultures

Human bone marrow-derived mesenchymal stromal cells (hBMSCs) were isolated from bone marrow aspirates of patients after informed consent, in accordance with a biobank protocol approved by the local Medical Ethics Committee (TCBio-08-001, University Medical Center Utrecht ^39^. hBMSCs were cultured in basal medium consisting of minimal essential medium (α-MEM; Gibco, 12571063), 10% heat-inactivated fetal bovine serum (FBS) (CAPRICORN, CP24-7059-HI), 0.2 mM L-ascorbic acid-2-phosphate (Sigma-Aldrich, 1713265-25-8), 100 U/mL penicillin with 100 mg/mL streptomycin (Gibco, 15140) and 1 ng/mL basic fibroblast growth factor (bFGF) (R&D Systems, 233-FB-500), which was added fresh every medium change. The basal medium was replaced twice per week. The cells were passaged when 80% confluency was reached using 0.25% Trypsin-EDTA (Gibco, 25200072). At passage 4, hBMSCs were used for subsequent experiments. Basal medium, supplemented with EV-free FBS (depleted of EVs by ultracentrifugation at 120,000 g for 16h), was added to the cells with/without microalgae EVs according to experimental design.

The chondrogenic cell line, ATDC5s, was cultured in basal medium composed of DMEM/F-12 (Gibco, 11320033), supplemented with 5% FBS and 100 U/mL penicillin with 100 µg/mL streptomycin (Gibco, 15140). For monolayer chondrogenic differentiation, 2.1 × 10⁵ cells per well were plated in a 48 well plate (Greiner) and cultured in chondrogenic medium consisting in high-glucose DMEM (Gibco, 31966), 1% ITS Premix (Corning, 354352), 40 µg/mL L-proline (Sigma-Aldrich, P5607), 10 ng/mL TGF-β (PeproTech, 100-21), 0.1 µM dexamethasone (Sigma-Aldrich, D4902), 50 µg/mL ascorbate 2-phosphate (Sigma-Aldrich, A8960), and 100 U/mL penicillin with 100 µg/mL streptomycin (Gibco, 15140). For pellet chondrogenic differentiation, 2 × 10⁵ of ATDC5 per well were plated in a 96 well U-bottom plates (Greiner, E22073N3), centrifuged at 300 g for 4 mins, and cultured in ATDC5 chondrogenic medium.

### 2.5 EV uptake assay

To visualize microalgal EVs uptake by mammalian cells, hBMSCs were treated with 20 µg/ml of CellMask™ (Invitrogen, C10046) labelled microalgae EVs as previously described ^40^. Briefly, EVs were incubated for 10 mins in dark conditions in CellMask™ solution (1:1000 in PBS). EVs were centrifuged at 120,000 g for 70 mins to remove the unbound dye using an Optima XE-90 ultracentrifuge with a SW 32 Ti rotor (Beckman Coulter, USA). Labelled-EVs were resuspended in PBS and added to hBMSCs, which seeded in 48 well plate and cultured overnight in expansion medium (containing EV-free FBS), and incubate for 16h at 37°C and 5% CO_2_. hBMSCs were washed twice with PBS, then fixed with 10 % formalin for 10 min at room temperature. After two washes with PBS, cells were permeabilized with 0.1% Triton X-100 for 10 min. After washing twice with PBS, samples were blocked with 2% BSA in PBS for 2 min. Blocking solution was removed and samples were incubated with 2.5 µg/mL of fluorescently conjugated phalloidin (in 2% BSA) for 20 min in the dark. After washing twice with PBS, nuclei were stained with 10 ng/mL of DAPI (in PBS) for 5 min in the dark, washed with PBS and images were acquired using confocal microscopy (Leica Microsystems, Germany). The excitation/emission wavelengths of DAPI, phalloidin and CellMask™ were 410/494 nm, 506/634 nm, and 669/754 nm, respectively.

### 2.6. hBMSCs proliferation and migration

The effects of microalgae EVs on hBMSCs proliferation was assessed via quantification of metabolic activity and DNA content. Briefly, hBMSCs (1 x 10^4^ cells/cm^2^) were treated with basal medium supplemented with or without 2 µg/mL of microalgae EVs and incubated for 1, 3 and 7 days. Cells cultured in basal medium was used as the control. At each time point, metabolic activity and DNA content was evaluated. To assess metabolic activity, the AlamarBlue reagent (44 mM of resazurin sodium salt, Sigma-Aldrich, R7017) was added and incubated for 4h at 37°C in the dark. Fluorescence was measured in a microplate reader (CLARIOstar Plus, BMG LABTECH, Ortenberg, Germany) using an excitation/emission of 530/590 nm. To assess DNA content, the Quant iT™ PicoGreen® dsDNA kit (Thermo Fisher Scientific, P11495) was used according to the manufacturer’s instructions. Briefly, cells were washed twice with PBS, then lysed with 0.1% Triton-X solution in PBS, followed by three cycles of freeze-thaw. Samples were diluted in TE buffer (10 mM Tris-HCl, 1 mM EDTA, pH 7.5) and transferred to a 96 well plate (100 µL per well), after which 100 µL of the PicoGreen working reagent was added and plates were incubated for 5 mins at room temperature in the dark. Fluorescence was measured using the mentioned microplate reader with an excitation/emission of 485/520 nm.

To assess migration rate, the scratch assay was performed as previously described ^41^. Briefly, cells were cultured in 6 well plates (30 × 10^3^ cells/cm^2^) in basal medium for 24h. A linear scratch was made using a sterile 20 µL pipette tip. After a PBS wash, basal medium (containing EV-depleted FBS) supplemented with/without EVs were added to the cells. Images were captured at day 0, 1 and 2 using an inverted microscope (DMi1, Leica Microsystems, Wetzlar, Germany). The scratch area was quantified using the software FIJI imageJ, the percentage of closure was calculated with the following formula [5], where *A0* is the area immediately after creating the scratch and *At* is the area at day 1.

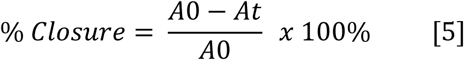

### 2.7. Cytokine-induced OA-like model

To evaluate the anti-catabolic effects of EV treatment, a cytokine-induced OA-like inflammatory model using the chondrogenic cell line ATDC5s was established. ATDC5s cultured in chondrogenic medium, as monolayers or as pellets, were exposed to either 10 ng/mL IL-1β (PeproTech, 200-01B) or 10 ng/mL TNF-α (PeproTech, 300-01A) ^42^. For the EV treated groups, ATDC5s were treated with 2 µg/mL of microalgae EVs. Cytokines and/or EVs were added fresh at each medium change.

Safranin-O staining was conducted to visualize cartilage-like glycosaminoglycan (GAG) formation. For chondrogenically differentiated ATDC5 monolayers, samples were stained with a modified Safranin-O protocol ^43^. Briefly, ATDC5 monolayers were washed twice with PBS and fixed with 10% formalin for 20 mins. After three washes with PBS, samples were incubated with 0.1% Safranin-O solution (Sigma Aldrich, 33209) for 30 mins. Following three washes with PBS, the samples were imaged using a stereoscopic microscope (Olympus SZ61, Olympus Corporation, Tokyo, Japan).

For ATDC5 chondrogenic pellets, samples were washed with PBS and placed in a drop of 5% low melt agarose (SeaKem®LE Agarose, Lonza), fixed in 10% formalin overnight and dehydrated by sequential immersion in: 70% ethanol (1h), 90% ethanol (1h), 96% ethanol (twice, 1h each), 100% ethanol (1h), 100% ethanol overnight and Xylene (twice, 1h each). Samples were then embedded in paraffin and sectioned in 5 µm using a rotatory microtome (HM 340E, Thermo Fisher Scientific, Waltham, MA, USA). Sections were deparaffinized and hydrated by sequential immersion in Xylene (twice, 5 min each), 100% ethanol (twice, 3 min each), 96% ethanol (twice, 3 min each), 70% ethanol (twice, 3 min each), and distilled water (three times for 2 min each). Safranin O/Fast Green staining was performed by placing the sections in Weigert’s hematoxylin for 5 min, washed under running tap water for 10 min and rinsed in distilled water. Sections were then counterstained with 0.4% aqueous Fast Green (MP Biomedicals, 0219517825) for 4 min. The slides were immersed in 0.125% Safranin O solution (Sigma-Aldrich, 33209) for 5 min after rinsing twice with 1% acetic acid (1 and 4 min respectively). Subsequently, slides were quickly rinse with 3 times 100% ETOH changes and transferred into Xylene for 5 min before mounting. Images were taken using a bright-field optical microscope (Olympus BX51, Olympus Corporation, Tokyo, Japan).

Dimethyl methylene blue (DMMB) assay was used to quantify GAG production. For monolayer samples, 100 µL of Milli-Q water was added to each well, then three freeze-thaw cycles were applied, and samples were subsequently incubated overnight at 60°C in 100 µL of papain solution (papain 250 µg/mL; cysteine HCl 1.57 mg/mL) prepared in 2× papain buffer (0.2 M sodium phosphate, 10 mM EDTA, pH 6.0). To quantify GAGs, papain-digested samples were diluted 1:30 in PBS-EDTA (40 mM Na₂HPO₄, 53.1 mM NaH₂PO₄·2H₂O, 10 mM disodium EDTA), transferred in duplicates of 50 µL per well to a 96 well plate and combined with 100 µL of DMMB solution (50 µM 1,9 dimethyl methylene blue, 40.6 mM NaCl, 40.5 mM glycine, 0.5% v/v ethanol, pH 3). The absorbance was measured at 525 and 595 nm wavelengths with previously mentioned microplate reader. For ATDC5 pellets, samples were first freeze-dried and then digested with above mentioned papain solution.

### 2.8. Statistical analysis

All statistical analyses were performed using GraphPad Prism 10 (GraphPad Software, San Diego, CA, USA). When comparing two groups, data was analysed using Student’s t-test after confirming for equality of variances with an F-test. For experiments involving more than two groups, one-way ANOVA was performed followed by multiple comparisons. In the case which involves two independent variables, two-way ANOVA were performed. All error bars are mean ± SD of at least three independent experiments. Statistically significant P values are indicated in figures and legends as *P ≤ 0.05, **P ≤ 0.01, ***P ≤ 0.001 and ****P ≤ 0.0001, ns represents no significant difference (P > 0.05).

## 3. Results

### 3.1. The Effects of Light Exposure Conditions on Microalgae Cultivation

To refine the manufacture of microalgae EVs, we initially determined the influence of the cultivation conditions on the viability and growth kinetics of different microalgae species. The cultivation conditions involved a photoperiod of either 0, 16, or 24h per day, for a total period of 2 weeks. Figure 2A shows the microscopic images of the different microalgae species assessed, showcasing their distinct morphological features. Macroscopic images highlight the impact of increased exposure of light irradiation on the intensity of pigmentation within the medium of different microalgae cultures following 2 weeks cultivation (Fig. 2B). A light exposure-dependent increase in microalgae number was observed in all species assessed, with *Leptolyngbya* exhibiting significantly increased microalgae number compared to *Chlorella* (4.8-fold), *CC1690* (7.3-fold) and *Synechococcus* (9-fold) (P ≤ 0.001) (Fig. 2C). A similar normalized microalgae biomass profile was also observed (Fig. S1). The impact of light irradiation on microalgae viability was assessed through live/dead staining and semi-quantification (Fig. 2D, E). Our results show that all microalgae species exhibited high viability (≥ 80%) after 2 weeks culture in different light irradiation periods, with *Chlorella* exhibiting significantly lower viability when compared to *Synechococcus*, *Leptolyngbya* and *CC1690* (P ≤ 0.0001). The 16h light irradiation exhibited the highest microalgae viability when compared to the 0h (P ≤ 0.01) and 24h light cycles (P > 0.05) after 2 weeks culture.

**Figure 2.**
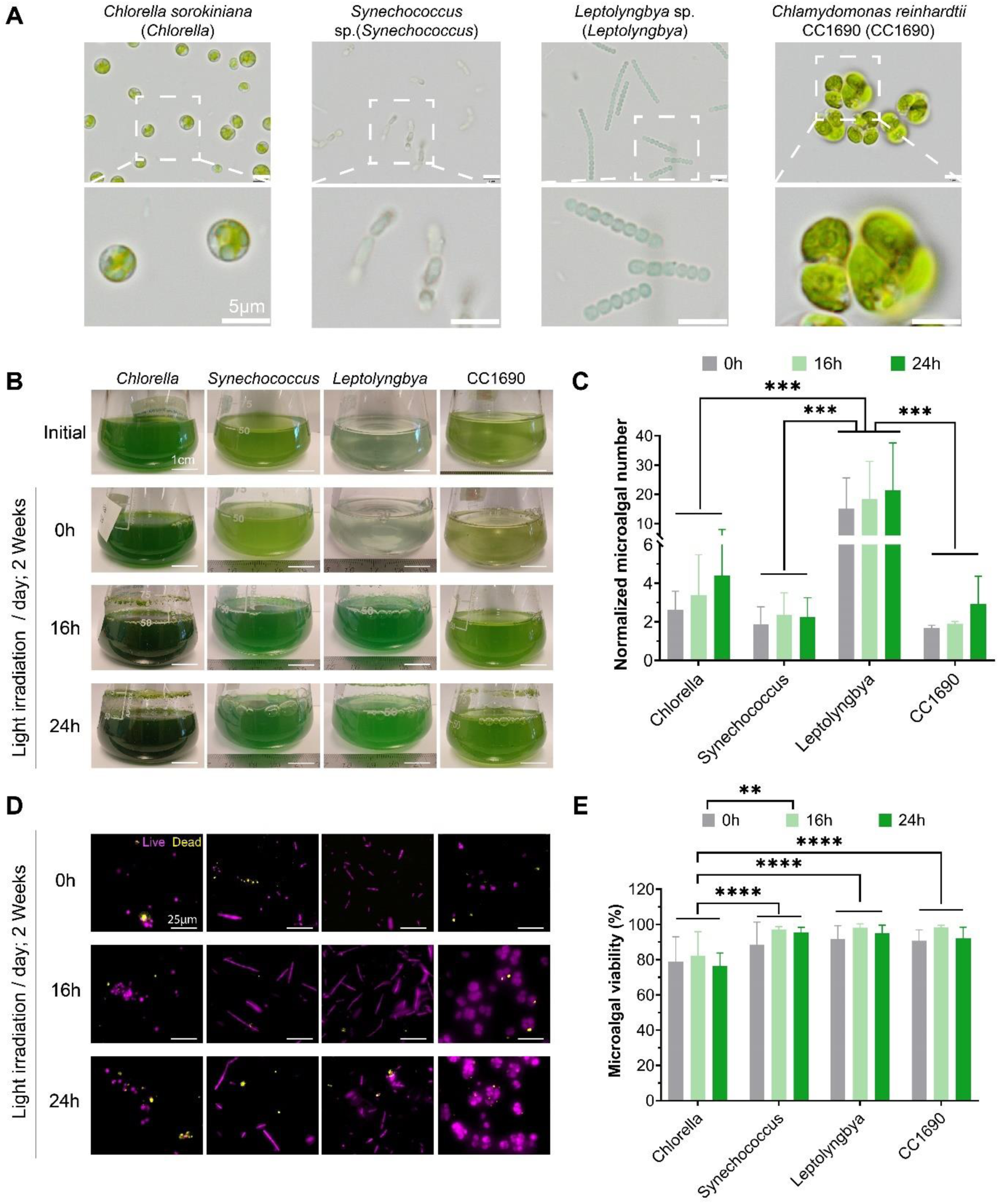
The effect of photoperiod on the growth and survival of different microalgae species. A) Microscopic images of different microalgae species. The effects of different cultivation conditions by 2 weeks, on B) microalgae medium pigmentation, C) normalized microalgae number, N_t_ (n = 3), D) live/dead fluorescent imaging of microalgae, E) microalgae viability (n=6). Data expressed as mean ± SD. **P ≤ 0.01, ***P ≤ 0.001 and ****P ≤ 0.0001.

### 3.2. The influence of Light Exposure Conditions on Microalgae EV production

We next evaluated the influence of microalgae light exposure conditions on EV production yield. Figure 3A provides an overview of the microalgae cultivation and EV isolation/purification used in this study. Our findings showed that all microalgae species were able to secrete EVs during the different cultivation conditions (Fig. 3B). *Leptolyngbya* exhibited the highest EV protein content, which was significantly higher when compared to *Synechococcus* (P ≤ 0.01), *Chlorella* (P ≤ 0.001), and *CC1690* (P ≤ 0.001). A light-dependent increase in EV protein content was observed for *Synechococcus and Leptolyngbya*, whilst light exposure negatively affected the production of EV protein per microalgae from *CC1690*. For *Chlorella*, light exposure did not affect EV protein content (Fig. 3B). In more detail, for *Synechococcus*, 24h light irradiation significantly increased EV protein content compared to 0h light exposure (P ≤ 0.05); for *Leptolyngbya*, both 16h (P ≤ 0.001) and 24h (P ≤ 0.001) exhibited significantly increased EV protein yields when compared to 0h light exposure; for *CC1690*, no effect on EV protein yields was observed while increasing light irradiation (P > 0.05).

**Figure 3.**
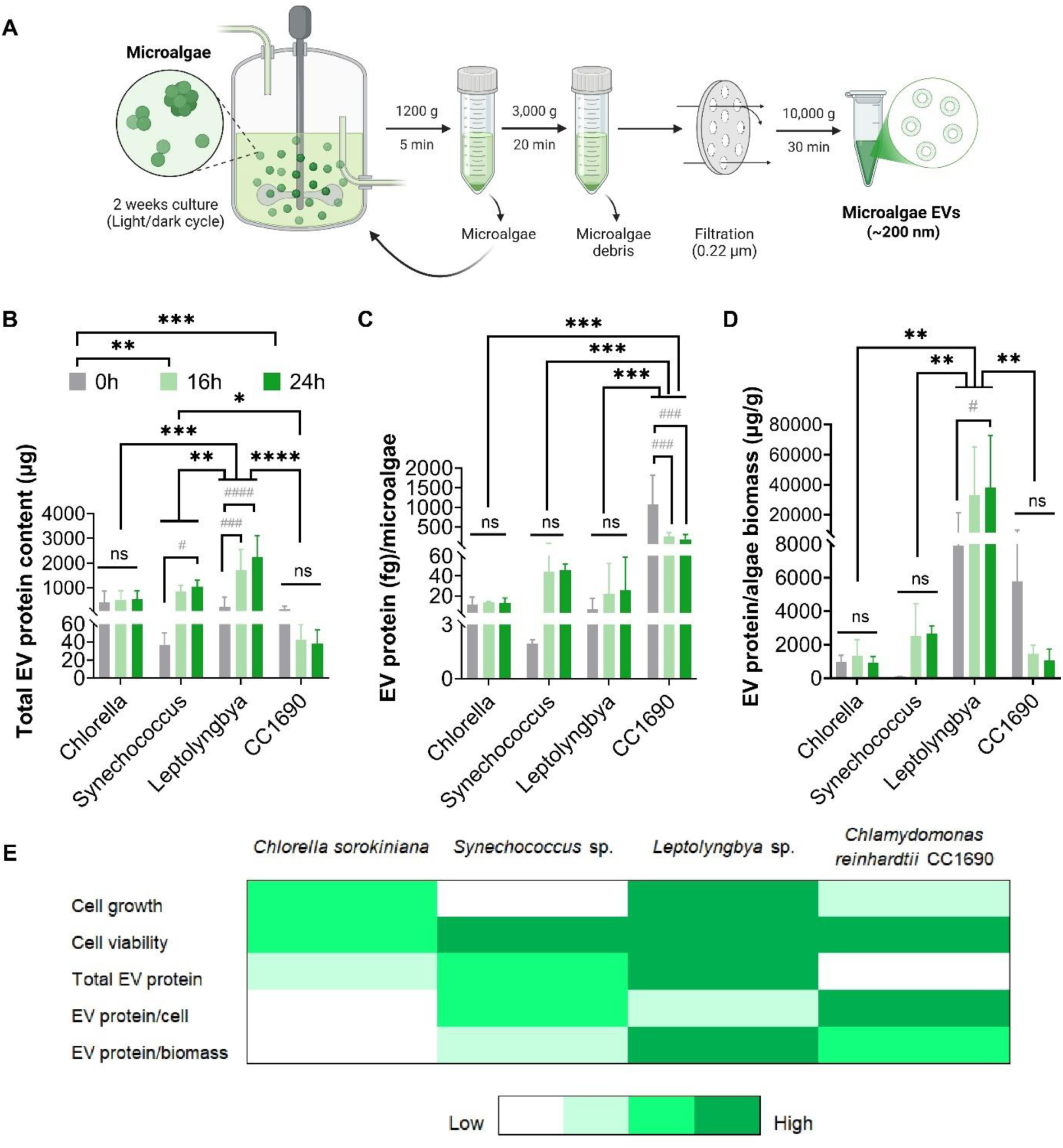
The effect of microalgae biomanufacturing conditions of EV production. A) Schematic overview of the microalgae cultivation and EV isolation procedure. Quantification of EV production by B) total EV protein content, C) EV protein per microalgae, D) EV protein per microalgae biomass. E) Summary of the key outputs regarding microalgae cultivation and EV production. Data expressed as mean ± SD. *P ≤ 0.05, **P ≤ 0.01, ***P ≤ 0.001 and ****P ≤ 0.0001. ^#^P ≤ 0.05, ^###^P ≤ 0.001, and ^####^P ≤ 0.0001 indicate significant differences among light conditions within the same microalgae species. ns = P > 0.05.

To determine the influence of cultivation conditions on the EV production per microalgae, EV protein content was normalised with the microalgae number (Fig. 3C) and microalgae biomass (Fig. 3D, biomass shown in Fig. S1). Following normalisation with microalgae number, *CC1690* exhibited the highest EV protein content, which was significantly higher than the *Synechococcus* (P ≤ 0.001), *Leptolyngbya* (P ≤ 0.001), and *Chlorella* (P ≤ 0.001) groups (Fig. 3C). Normalisation with microalgae biomass, *Leptolyngbya* displayed the highest EV protein yield, followed by *CC1690* (P ≤ 0.01), *Synechococcus* (P ≤ 0.01), and *Chlorella* (P ≤ 0.01) (Fig. 3D).The influence of different light irradiation cycles on the EV production yield within each microalgae species remains consistent to the results shown in Figure 3B. Figure 3E provides an overview of important outputs influencing the selection of an optimum microalgae EV factory, such as microalgae growth kinetics, viability, and EV production yields. *Leptolyngbya* exhibited high algae growth kinetics, whilst *Leptolyngbya*, *Synechococcus* and *CC1690* displayed high cell viability. Total EV production yield was highest in the *Leptolyngbya* group, where normalization with cell number and biomass, CC1690 and *Leptolyngbya*/*CC1690* exhibited the highest EV protein values, respectively. Taken together, *Leptolyngbya* EVs were selected for further analysis due to the increased growth kinetics, viability and EV production yields.

### 3.3. Characterisation of *Leptolyngbya-*derived EVs

Following the selection of *Leptolyngbya* as our microalgae EV source, we further explored refining our manufacturing conditions to improve EV production yields. Due to the importance of light exposure to *Leptolyngbya* growth kinetics, we initially assessed the influence of light irradiation cycles (16 and 24h) on the antioxidant capacity of isolated EVs through the DPPH• scavenging assay. Our findings showed that EVs produced from the 16h light irradiation cycle exhibited a 1.59-fold increase in the DPPH• scavenging activity (15%) when compared to the 24h group (9.44%) (P ≤ 0.05) (Fig. 4A). The influence of microalgae medium collection frequency on the EV production yield was next evaluated. The quantity of EVs produced from the “Standard” cultivation protocol (1 collection after 2-week cultivation) and a “Frequent” cultivation protocol (4 collections during 2-week cultivation) was assessed. Our findings showed that when compared to the “Standard” cultivation protocol used for microalgae cultivation, our “Frequent” harvesting regimen substantially increased vesicles production yield regarding EV protein content and particle number by 2.64-fold and 270-fold, respectively (Fig. S2). The 16h light irradiation cycle, in conjunction with the “Frequent” harvesting regimen were employed for the rest of this study. Following the isolation of EVs, TEM imaging showed the isolated nanoparticles exhibited a typical spherical morphology with a heterogenous size ranging from ∼50 - 300 nm (Fig. 4B). NTA analysis showcased that the isolated Lepto-EVs exhibited an average diameter of 161.7 ± 3.0 nm and a total concentration of 1.21 x 10^11^ particles/ml (Fig. 4C). To confirm the isolated nanoparticles were EVs, immunoblotting was conducted using the Exo-Check™ Exosomes Antibody Array. Immunoblotting confirmed the presences of EV-associated markers (CD63, EpCAM, ANXA5, TSG101, FLOT1, ICAM, ALIX, and CD81) (Fig. 4D).

**Figure 4.**
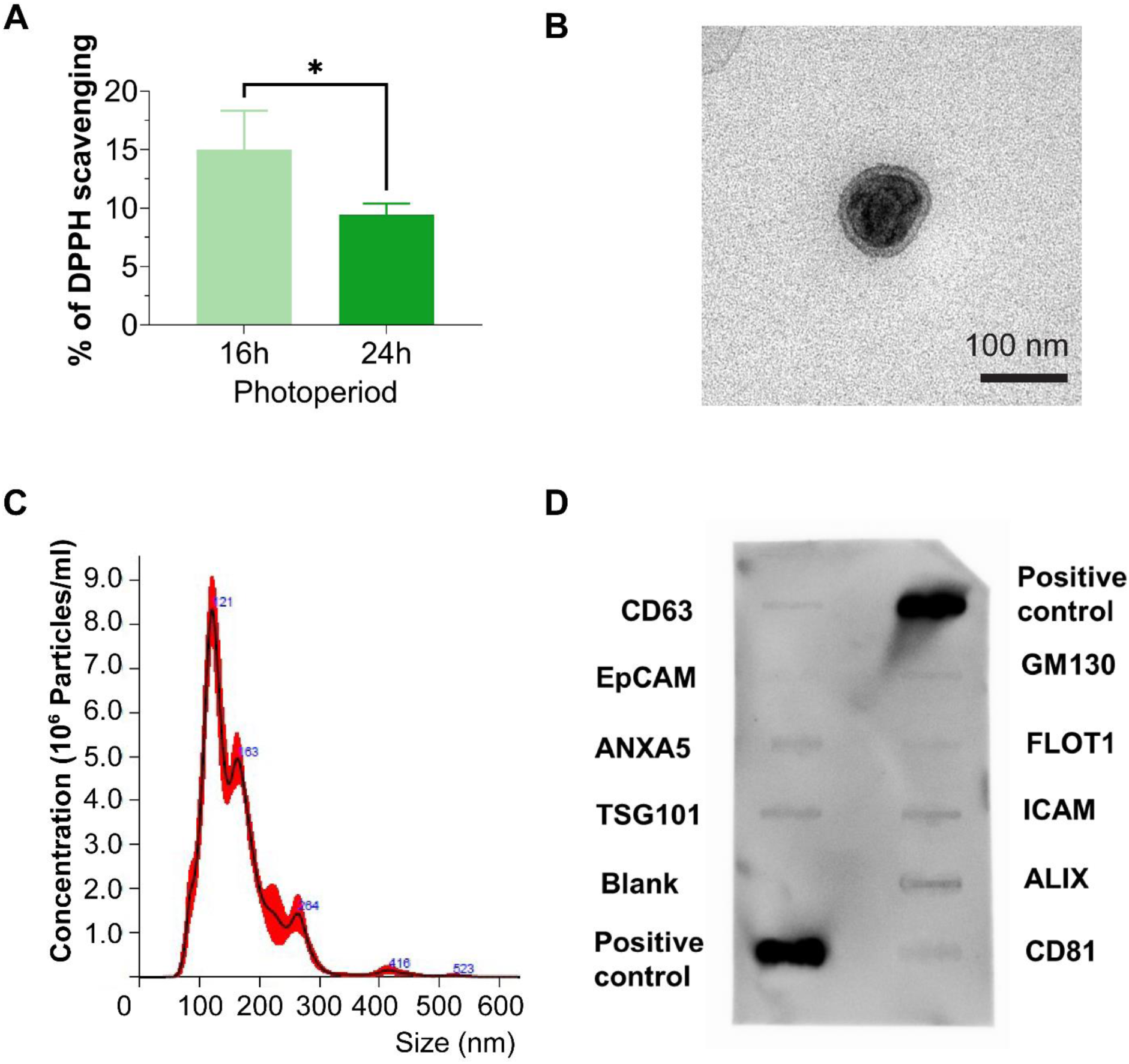
Characterization of *Leptolyngbya*-derived EVs. A) Percentage of DPPH scavenging capacity of EVs cultured in different light/dark cycles (n=3) B) TEM, C) NTA, and D) Quality assessment of isolated EVs with the Exo-Check™ antibody array. Lepto-EVs were assayed to detect eight EV-associated protein markers, including CD63, ANXA5 (annexin A5), CD81, FLOT1 (flotilin-1), ICAM1 (intercellular adhesion molecule 1), ALIX (programmed cell death 6 interacting protein), TSG101 (tumour susceptibility gene 101), EpCAM (epithelial cell adhesion molecule), and a control for cellular contamination, GM130 (cis-golgi matrix protein). Data expressed as mean ± SD. *P ≤ 0.05.

### 3.4. Lepto-EVs Promote the Proliferation and Migration of hBMSCs

The influence of Lepto-EVs on hBMSC general behaviour was initially assessed by evaluating their uptake into recipient cells. CellMask™-labelled Lepto-EVs were successfully internalized by hBMSCs, with the labelled EVs located within the cytoplasmic regions of the cell (Fig. 5A). Treatment with Lepto-EVs significantly increased the hBMSC proliferative capacity over a 7-day period when compared to the untreated cells, assessed by quantifying metabolic activity and DNA content (Fig. 5B, C) (P ≤ 0.01 - 0.0001). Additionally, hBMSC migration was substantially accelerated following Lepto-EV treatment, resulting in a 1.44-fold increase in % gap closure (58.72%) when compared to the untreated cells (40.70%) at 1 day (Fig. 5D, E) (P ≤ 0.05).

**Figure 5.**
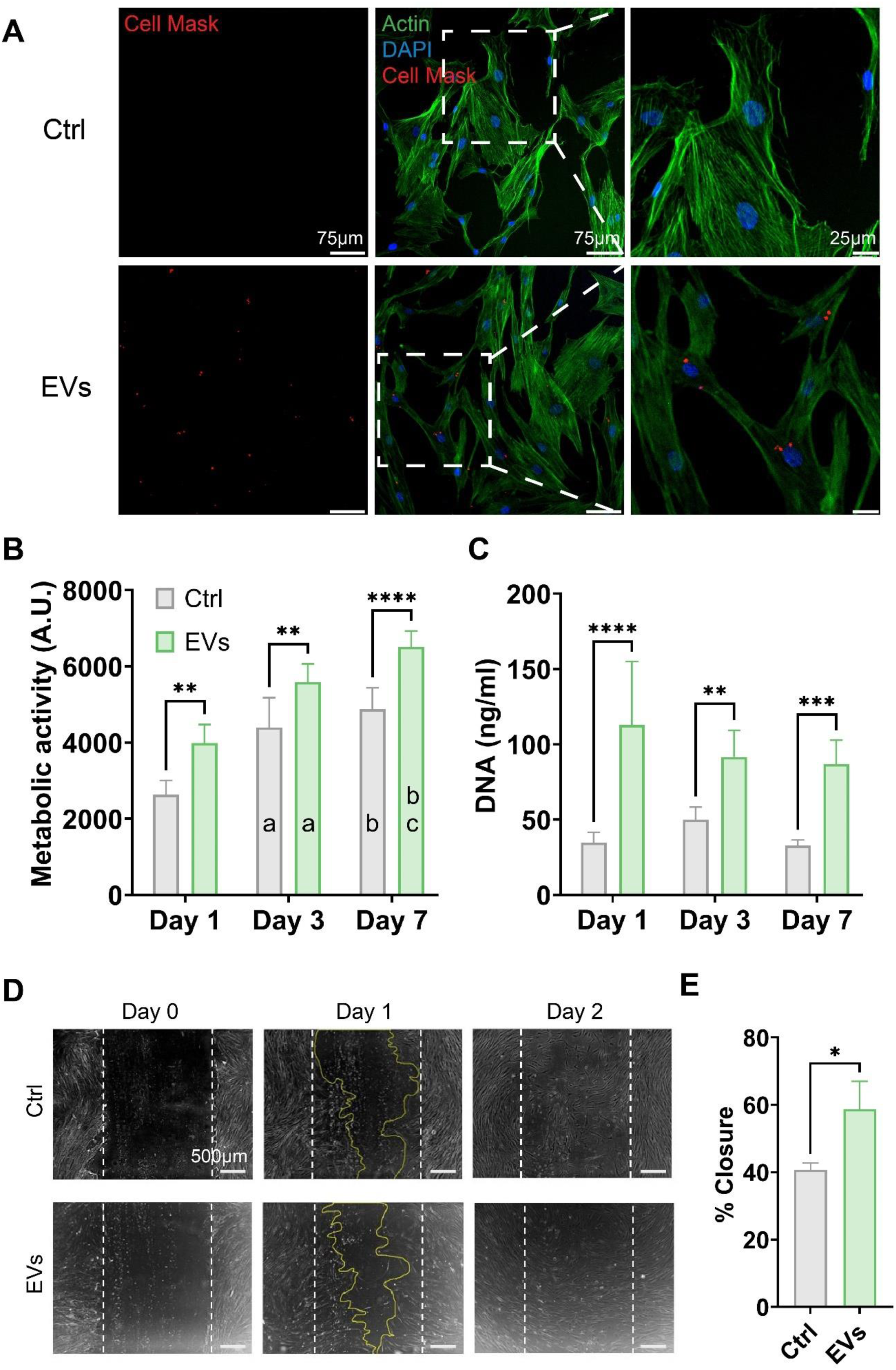
The influence of Lepto-EVs on hBMSCs general behaviour. A) Fluorescent images of Cell Mask-labelled Lepto-EV uptake by hBMSCs. The effects of Lepto-EVs on hBMSCs B) metabolic activity, C) DNA content, and D) migration microscopic images, E) quantified % gap closure at 1 day. Data expressed as mean ± SD. *P ≤ 0.05, **P ≤ 0.01, ***P ≤ 0.001 and ****P ≤ 0.0001. Letters (a–c) indicate significant differences within the same group (at least *P ≤ 0.05) between (a) Day 1 and Day 3, (b) Day 1 and Day 7, and (c) Day 3 and Day 7, respectively.

### 3.5. Therapeutic Effects of Lepto-EVs within a Cytokine-Induced OA-like Model

The effects of Lepto-EV treatment on alleviating cartilage-like matrix degradation were assessed using a cytokine-induced OA-like model. Within the 2D model, positive staining for GAGs was observed in all treatment groups (Fig. 6A). TNF-α and IL-1β-treated groups exhibited reduced GAG staining intensity when compared to the untreated control at day 3. The Lepto-EV-treated cytokine groups exhibited a similar degree of GAG staining when compared to the untreated control. To further investigate the influence of Lepto-EV treatment on alleviating cytokine-induced cartilage-like matrix degradation, the quantification of GAG content was conducted (Fig. 6B). Our findings showed a reduction in GAG content following TNF-α and IL-1β treatment when compared to the untreated control at both time points. Lepto-EV treatment did not improve GAG production in both the TNF-α and IL-1β treated groups at day 1 (P > 0.05), but improved GAG production at day 3 (P ≤ 0.001 - 0.0001) when compared to the EV-free cytokine treated cells. An ATDC5-based 3D *in vitro* model ^44,45^ was employed to further evaluate the influence of Lepto-EV treatment on reducing cytokine-induced matrix degradation within a more physiologically relevant system. Safranin-O staining was conducted to qualitatively assess GAG production. TNF-α and IL-1β treatment resulted in reduced Safranin-O staining intensity when compared to the cytokine-free control after 14 days of culture (Fig. 6C). Lepto-EV treated cytokine groups exhibited increased Safranin-O staining intensity when compared to the EV-free cytokine-treated groups. Following the quantification of GAG content, our findings showed that Lepto-EV treatment increased GAG production in the TNF-α and IL-1β-treated, as well as in the cytokine-free groups (Fig. 6D).

**Figure 6.**
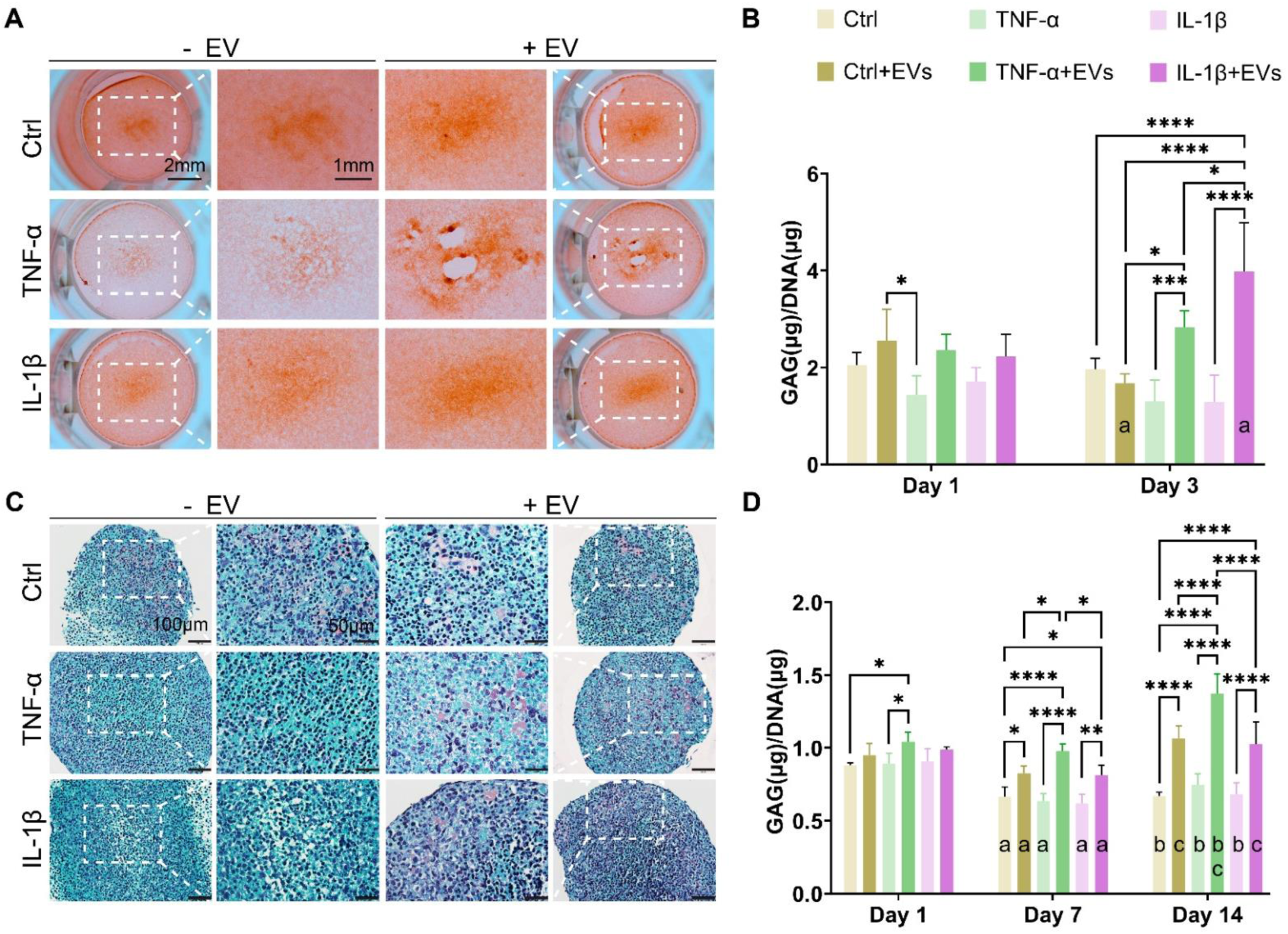
The effects of Lepto-EV treatment within a cytokine-induced OA-like *in vitro* model. A) Safranin-O staining and B) GAG content normalized by DNA content within a cytokine-induced 2D ATDC5 OA-like model (n ≥ 3). a) at least *p ≤ 0.05 between Day 1 and Day 3 within the same group. C) Safranin O staining and D) GAG content normalized by DNA content within a cytokine-induced 3D ATDC5 OA-like model (n=5). Letters (a–c) indicate significant differences within the same group (at least *P ≤ 0.05) between (a) Day 1 and Day 7, (b) Day 1 and Day 14, (c) Day 7 and Day 14, respectively. Data are expressed as mean ± SD. *P ≤ 0.05, **P ≤ 0.01, ***P ≤ 0.001 and ****P ≤ 0.0001.

## 4. Discussion

Despite the growing interest in EVs as precision nanotherapeutics, their clinical translation remains constrained by fundamental limitations in mammalian cell-based production systems, including donor variability, low yield, labour-intensive isolation, and scalability challenges ^46,47^. Here, we propose an alternative strategy using microalgae-derived EVs, leveraging these photosynthetic microorganisms’ unique capacity for scalable and economical production ^34,48^. We performed a first systematic optimization of microalgal EV production for potential therapeutic use, focusing on a clinically relevant OA disease model. Through comparative selection and iterative refinement of cultivation parameters, *Leptolyngbya* emerged as a lead species, demonstrating superior growth kinetics, cell viability, and EV productivity under tailored photoperiod and harvesting conditions. Importantly, Lepto-EVs exhibited potent antioxidant, and anti-catabolic effects, underscoring their potential as the next-generation, cost-effective and sustainable EV-based nanotherapeutic for OA.

Our investigations into the biomanufacturing of microalgae EVs demonstrate the profound influence of cultivation conditions, particularly light exposure, on both algal growth and subsequent EV secretion. Our findings indicate that both microalgae growth and subsequent EV production are species- and light-dependent processes. For instance, *Leptolyngbya* demonstrated a clear advantage, showing superior growth kinetics and viability across different photoperiods, which directly correlated with a significantly higher total EV protein yield. This suggests that optimized growth conditions are a prerequisite for efficient EV production. In contrast, while *Chlorella* showed increased EV protein content with more light, its overall relatively lower viability and growth compared to other species present a challenge for its large-scale EV production efficiency. The unique response of *CC1690*, where increased light negatively impacted EV production despite high viability, underscores that the relationship between cellular health, growth, and EV secretion is not always linear and is highly species-specific. The normalization of EV yield to microalgae number and biomass further refines this understanding, revealing that while *Leptolyngbya* is a top producer on a per-culture basis, *CC1690* may be a highly efficient producer on a per-cell basis. Beyond influencing microalgae growth and EV yield, our findings showcased that light exposure also plays a pivotal role in shaping the antioxidant capacity of Lepto-EVs. While continuous (24h) illumination supported the highest growth rate and EV production, a 16h light regimen markedly enhanced the antioxidant properties of the vesicles. Excessive light intensity likely compromises microalgae’s antioxidant capacity by inducing photoinhibition and excessive reactive oxygen species (ROS) generation, which can overwhelm cellular defence systems and degrade or suppress the synthesis of antioxidant molecules ^49,50^. Therefore, optimizing the light regime represents an effective strategy to fine-tune the therapeutic potential of Lepto-EVs, particularly for mitigating the excessive ROS associated with OA pathology.

In addition to light exposure, our results demonstrate the significant impact of medium harvesting frequency on EV production. Frequent medium replacement in microalgae cultures sustains nutrient availability, removes inhibitory metabolites, and maintains stable pH and gas conditions, keeping algae in a metabolically active state ^51^. This promotes faster growth and can enhance EV production, as vesicle release is linked to both cellular metabolism and transient stress responses ^52^. By preventing nutrient limitation and self-inhibition while providing occasional mild stress cues, medium exchange effectively supports higher biomass accumulation and increased vesicle secretion. This approach contrasts with studies in the literature which typically relied on a single collection ^53,54^, highlighting that our frequent collection strategy may contribute to sustained EV yields and more dynamic insights into vesicle production over time. These results are crucial for establishing a rational framework for microalgae-based EV manufacturing, confirming that selecting the appropriate species, optimizing light cycles and refining harvesting frequency are fundamental steps for maximizing the therapeutic potential of these nanovesicles.

Biophysical characterization confirmed that the obtained Lepto-EVs possess key hallmarks indicative of EVs, including their nanoscale size, spherical morphology, and expression of classical EV-associated protein markers, such as CD63, EpCAM, ANXA5, TSG101, FLOT1, ICAM, ALIX, and CD81, consistent with previous findings in the literature ^53–55^. Our findings showed that these Lepto-EVs were effectively internalized into the cytoplasm of recipient hBMSCs, demonstrating the inter-kingdom communication or cross-kingdom interaction between these plant-derived nanoparticles and human cells. Similar studies have demonstrated the internalization of EVs derived from *Spirulina platensis* (SP) ^56^ and *Tetraselmis chuii* ^54^ in ATDC5 and MDA-MB 231 cell lines, respectively.

The effective recruitment of endogenous cells is critical for subsequent tissue healing ^57,58^. Several studies have reported the capacity of EVs in stimulating the recruitment of progenitor cells in musculoskeletal applications ^21,59,60^. Functionally, our findings showed that the Lepto-EVs were not only cytocompatible, but significantly enhanced the proliferation and migration capacity of progenitor cells, crucial cellular processes conducive to cartilage repair. This is consistent with evidence in the literature demonstrating the cytocompatibility of microalgae-derived EVs, although this has primarily been assessed with cell lines ^53,54,56^. In recent years, there has been increasing studies investigating the xenogenic administration of human-, milk- and plant-derived EVs ^61–63^, particularly regarding their biosafety and biocompatibility. For instance, Adamo *et al*. administrated *Tetraselmis chuii*-derived EVs intravenously into immune-competent BALB/c mice and did not observe any noticeable local and systemic toxicity ^54^. Similarly, Liang *et al*. reported that weekly intra-articular injections of SP-EVs in an OA mouse model caused no detectable adverse effects ^56^. Our findings demonstrate the successful manufacture and isolation of Lepto-EVs, as well as their ability to interact and modulate the biological function of human-derived cells. These results underscore the potential of Lepto-EVs as a biologically active and biocompatible platform. Nonetheless, comprehensive safety assessments remain essential. Future studies should evaluate Lepto-EVs tolerability within *in vivo* studies, alongside detailed biodistribution information to fully define their therapeutic potential and translational applications.

To investigate the anti-catabolic effects of Lepto-EVs, a cytokine-induced OA-like *in vitro* model was employed as it provides a controlled and reproducible system to mimic the inflammatory and catabolic milieu of OA. This approach reduces donor variability, minimizes ethical and cost constraints, and enables high-throughput assessment of EV bioactivity compared with *ex vivo* explants or *in vivo* models ^64^. As such, it serves as an efficient first-line screening platform prior to validation in more complex tissue or animal models. The cytokine-dependent reduction in GAG content during chondrogenic differentiation observed in both 2D and 3D models, was consistent with findings of OA *in vitro* models in the literature ^65,66^. The findings from both 2D and 3D *in vitro* models provide evidence for the protective role of Lepto-EVs in mitigating cartilage-like matrix degradation associated with OA. In the 2D model, the reduction in Safranin-O staining and total GAG content following TNF-α and IL-1β stimulation was significantly countered by Lepto-EV treatment, with the GAG content returning to levels comparable to the untreated control. This suggests that Lepto-EVs can directly protect the cartilage-like extracellular matrix from inflammation-induced damage. The anti-catabolic effects of Lepto-EVs in our 2D ATDC5 model, is consistent with the findings from Liang *et al.,* who observed increased Sox9 and Col2a gene expression following SP-EV treatment of TNF-α stimulated ATDC5s ^56^. The results from the 3D model, which more closely mimics the physiological environment of cartilage ^67^, further support this conclusion. The marked reduction in GAG content observed in the cytokine-treated groups was substantially reversed by Lepto-EVs, demonstrating their ability to preserve the integrity of the cartilage-like extracellular matrix, a crucial aspect of OA therapy ^68,69^. The observed increase in GAG content in cytokine-stimulated cells treated with Lepto-EVs highlights a reparative effect, suggesting that these vesicles may not only prevent matrix degradation, but may also promote anabolism within these chondrogenic cells. Although ATDC5s present a useful chondrogenic cell line ^70,71^, it has been reported that the cartilage-like matrix produced from these cells is inherently less complex than that found in native articular cartilage ^72^. Moreover, the high proliferative capacity of these cells rather than the stable, low-proliferative nature of native chondrocytes ^73,74^, may overestimate effects on proliferative pathways while underrepresenting mechanisms relevant to quiescent or senescent chondrocytes. Nevertheless, the anti-catabolic findings observed in this study are likely mediated through the delivery of antioxidant and anti-inflammatory molecules inherent to the microalgal origin of the EVs, including phycocyanin, carotenoids, and phenolic compounds ^56,75,76^.

Our findings position Lepto-EVs as a potential next-generation cell-free nanotherapy for OA, with distinct advantages over existing biological approaches. While mammalian EVs can also carry bioactive cargo, their content is highly variable and donor-dependent ^77,78^. In contrast, microalgal EV composition can be more precisely modulated through environmental control, enabling a more standardized therapeutic product. Moreover, their environmental sustainability and ease of scale-up address key bottlenecks in the biomanufacturing pipeline, aligning with global priorities for green biotechnologies ^34,48^. However, several questions remain. The precise molecular cargo of Lepto-EVs, particularly their RNA and protein content, requires further elucidation to understand the mechanisms underlying their therapeutic effects. Additionally, while *in vitro* models provide important proof-of-concept data, *in vivo* studies are important to evaluate biodistribution, pharmacokinetics, immunogenicity, and long-term efficacy. Importantly, regulatory pathways for algae-derived nanotherapeutics remain underdeveloped, necessitating the dialogue with policymakers and standardization bodies. The clinical translation of plant- and microalgae-derived EVs remains in its early stages, largely due to regulatory and classification challenges. For pharmaceutical applications, mammalian-based EV products may be classified as biological medicinal products, requiring precise definition of active substances and elucidation of mechanisms of action ^79^. In contrast, plant-derived EV-based products may fall under the category of botanical drugs, for which demonstrating safety and efficacy is sufficient under FDA guidance ^80^. In the EU, plant-derived EV products could be regulated as advanced therapy medicinal products (ATMPs), necessitating thorough evaluation of pharmacological properties, manufacturing processes, and quality control. Comprehensive physicochemical profiling, safety validation, and standardized manufacturing are therefore essential to meet these regulatory demands and advance the clinical and commercial translation of plant-derived nanovesicles.

In addition to demonstrating potent regenerative effects *in vitro*, the present study underscores the economic and biomanufacturing potential of microalgae-derived EVs as next-generation nanotherapeutics. Unlike MSC-derived EVs, which are hampered by costly xeno-free media, and limited scalability ^81^, microalgal platforms can be cultivated in photobioreactors with minimal nutrient inputs and reliance on light and CO₂, thereby markedly reducing upstream costs and reducing the high carbon footprint associated with mammalian cell manufacture ^82,83^. Although downstream purification and regulatory frameworks remain to be fully optimized, the combination of rapid biomass accumulation, high EV yield, and sustainable (lower carbon footprint) production positions Lepto-EVs as a viable alternative to EVs derived from mammalian sources. By bridging nanomedicine, environmental biotechnology, and regenerative medicine, our findings advance microalgae EVs as a scalable, cost-efficient, and environmentally sustainable nanotherapeutic platform with significant translational potential in regenerative medicine.

## 5. Conclusion

In conclusion, this research underscores the critical need to refine microalgae biomanufacturing to enable the scalable production of EVs with clinical potential. Our comprehensive species selection successfully identified Lepto-EVs as a potent, cell-free, and environmentally sustainable nanotherapeutic platform. These vesicles demonstrated significant potential, specifically exhibiting anti-catabolic effects relevant to the treatment of OA. This foundational work is an important step in the establishment of a clear translational pathway for developing next-generation, algae-derived nanotherapeutics.

## Acknowledgements

1. M. W would like to acknowledge the China Scholarship Council (CSC202108440024) for the financial support. The authors would like to acknowledge Fengzheng Gao and Maria Barbosa at Wageningen University & Research for assistance with the calculation of microalgae cell number and biomass. Figures were created with BioRender.com.

## Author contributions

M.W and K.M - conceptualization. M.W and S.G - Investigation. M.W and S.G - Formal analysis. K.M - Project administration. J.M, K.M - Supervision. M.W - Writing – original draft. S.G, A.K.O, J.L.R, M.D.R, D.G, J.M, K.M - Writing – review and editing. All authors have read and agreed to the published version of the manuscript.

## Supplementary

**Supplementary Table 1:** Microalgal cultivation media.

|  | Artificial Seawater Medium |  |  | Chlorella Medium |  |  | Modified TAP medium |  |  |
| --- | --- | --- | --- | --- | --- | --- | --- | --- | --- |
|  | g/L | MW | mM | g/L | MW | mM | g/L | MW | mM |
| **Trace Element** |  |  |  |  |  |  |  |  |  |
| Na<sub>2</sub>EDTA·2H<sub>2</sub>O | 4.50E-02 | 372.24 | 1.20E-01 | 3.72E-02 | 372.24 | 1.00E-01 | 9.31E-03 | 372.24 | 2.50E-02 |
| MnCl<sub>2</sub>·2H<sub>2</sub>O | 1.71E-03 | 143.86 | 1.20E-02 | 1.30E-02 | 143.86 | 6.60E-02 | 1.19E-03 | 143.86 | 8.00E-03 |
| ZnSO<sub>4</sub>·7H<sub>2</sub>O | 6.60E-04 | 287.56 | 2.00E-03 | 3.20E-03 | 287.56 | 1.10E-02 | 7.20E-04 | 287.56 | 3.00E-03 |
| Co(NO<sub>3</sub>)<sub>2</sub>·6H<sub>2</sub>O | 7.00E-05 | 291.03 | 2.41E-04 |  |  |  |  |  |  |
| CuSO<sub>4</sub>·5H<sub>2</sub>O | 2.40E-05 | 249.69 | 9.61E-05 | 1.83E-03 | 249.69 | 7.00E-03 | 3.40E-04 | 170.48 | 2.00E-03 |
| CuCl<sub>2</sub>·2H<sub>2</sub>O |  |  |  |  |  |  | 3.50E-05 | 1235.86 | 2.85E-05 |
| Na<sub>2</sub>MoO<sub>4</sub>·2H<sub>2</sub>O | 2.42E-04 | 241.95 | 1.00E-03 |  |  |  |  |  |  |
| H<sub>2</sub>SeO<sub>3</sub> | 1.29E-05 | 128.97 | 1.00E-04 |  |  |  |  |  |  |
| Na<sub>2</sub>SeO<sub>3</sub> |  |  |  |  |  |  | 1.70E-05 | 172.95 | 9.95E-05 |
| NiSO<sub>4</sub>·6H<sub>2</sub>O | 2.63E-05 | 262.85 | 1.00E-04 |  |  |  |  |  |  |
| Na<sub>3</sub>VO<sub>4</sub> | 1.84E-05 | 183.91 | 1.00E-04 |  |  |  |  |  |  |
| K<sub>2</sub>CrO<sub>4</sub> | 1.94E-05 | 194.19 | 1.00E-04 |  |  |  |  |  |  |
| **Iron stock** |  |  |  |  |  |  |  |  |  |
| NaFeEDTA | 3.96E-02 | 367.05 | 1.08E-01 | 1.16E-01 | 367.05 | 3.16E-01 |  |  |  |
| FeCl<sub>3</sub>·6H<sub>2</sub>O |  |  |  |  |  |  | 5.40E-03 | 270.29 | 2.00E-02 |
| **Nitrate and Phosphate Source** |  |  |  |  |  |  |  |  |  |
| NaNO<sub>3</sub> | 2.00E+00 | 84.99 | 2.35E+01 | 2.00E+00 | 84.99 | 2.35E+01 | 2.00E+00 | 84.99 | 2.35E+01 |
| KH<sub>2</sub>PO<sub>4</sub> | 2.30E-01 | 136.09 | 1.69E+00 | 7.40E-01 | 136.09 | 5.44E+00 | 1.08E-01 | 136.09 | 7.94E-01 |
| Na<sub>2</sub>HPO<sub>4</sub>·2H<sub>2</sub>O |  |  |  | 2.60E-01 | 177.99 | 1.46E+00 | 5.40E-02 | 177.99 | 3.03E-01 |
| Na<sub>2</sub>EDTA | 6.00E-02 | 372.24 | 1.61E-01 |  |  |  |  |  |  |
| **Macronutrients** |  |  |  |  |  |  |  |  |  |
| NaCl | 2.45E+01 | 419.23 |  |  |  |  |  |  |  |
| Na<sub>2</sub>SO<sub>4</sub> | 3.20E+00 | 142.04 |  |  |  |  |  |  |  |
| CaCl<sub>2</sub>·2H<sub>2</sub>O | 8.00E-01 | 147.01 | 5.42E+00 | 3.00E-02 | 147.01 | 1.90E-02 | 5.00E-02 | 147.01 | 2.50E-02 |
| K<sub>2</sub>SO<sub>4</sub> | 8.50E-01 | 174.26 | 4.88E+00 |  |  |  |  |  |  |
| MgCl<sub>2</sub>·6H<sub>2</sub>O | 9.80E+00 |  | 4.82E+01 |  |  |  |  |  |  |
| MgSO<sub>4</sub>·7H<sub>2</sub>O |  |  |  | 4.00E-01 | 246.47 | 1.62E+00 | 1.00E-01 | 246.47 | 2.50E-02 |
| H<sub>3</sub>BO<sub>3</sub> |  |  |  | 6.10E-05 | 61.83 | 1.00E-03 |  |  |  |

**Supplementary Table 2:** Formula used for calculating microalgae cell number. DC: dilute coefficient.

| Microalgal species | Cell counting ($\times 10^6$ cells $\text{ml}^{-1}$) | $R^2$ |
| --- | --- | --- |
| *Chlorella sorokiniana* | $18.181 \times \text{OD750} \times 2.9 \times \text{DC} + 58.552$ | 0.9367 |
| *Synechococcus* sp. | $107.020 \times \text{OD750} \times 2.74 \times \text{DC} + 8.049$ | 0.9132 |
| *Leptolyngbya* sp. | $(27.421 \times \text{OD750} \times 2.21 \times \text{DC} \times 100 - 59.365) / 100$ | 0.8713 |
| *Chlamydomonas reinhardtii* CC1690 | $4.008 \times \text{OD750} \times 1.29 \times \text{DC} + 1.407$ | 0.953 |

**Supplementary Table 3:** Formula used for calculating microalgal dry biomass. DC: dilute coefficient.

| Microalgal species | Biomass counting ($\times \text{g L}^{-1}$) | $R^2$ |
| --- | --- | --- |
| *Chlorella sorokiniana* | $0.3354 \times \text{OD750} \times 2.9 \times \text{DC} - 0.3868$ | 0.937 |
| *Synechococcus* sp. | $0.334 \times \text{OD750} \times 2.74 \times \text{DC} + 0.2087$ | 0.913 |
| *Leptolyngbya* sp. | $(0.4672 \times \text{OD750} \times 2.21 \times \text{DC} - 0.2422)$ | 0.956 |
| *Chlamydomonas reinhardtii* CC1690 | $0.657 \times \text{OD750} \times 1.29 \times \text{DC} + 0.031$ | 0.99 |

**Supplementary Figure 1.**
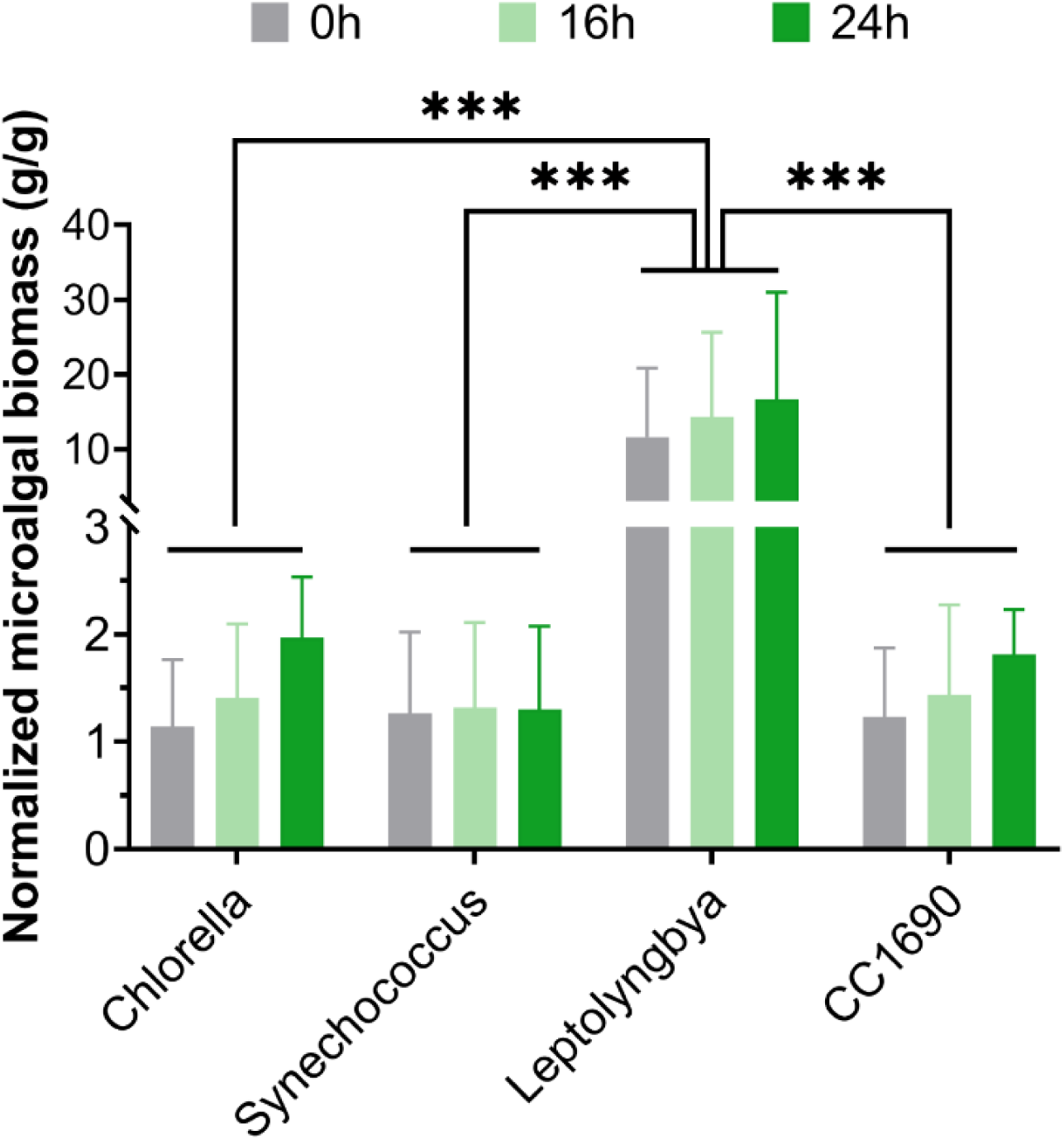
Normalized microalgal biomass for four algae species exposed to different light/dark regimens. Data are expressed as mean ± SD. ***P ≤ 0.001.

**Supplementary Figure 2.**
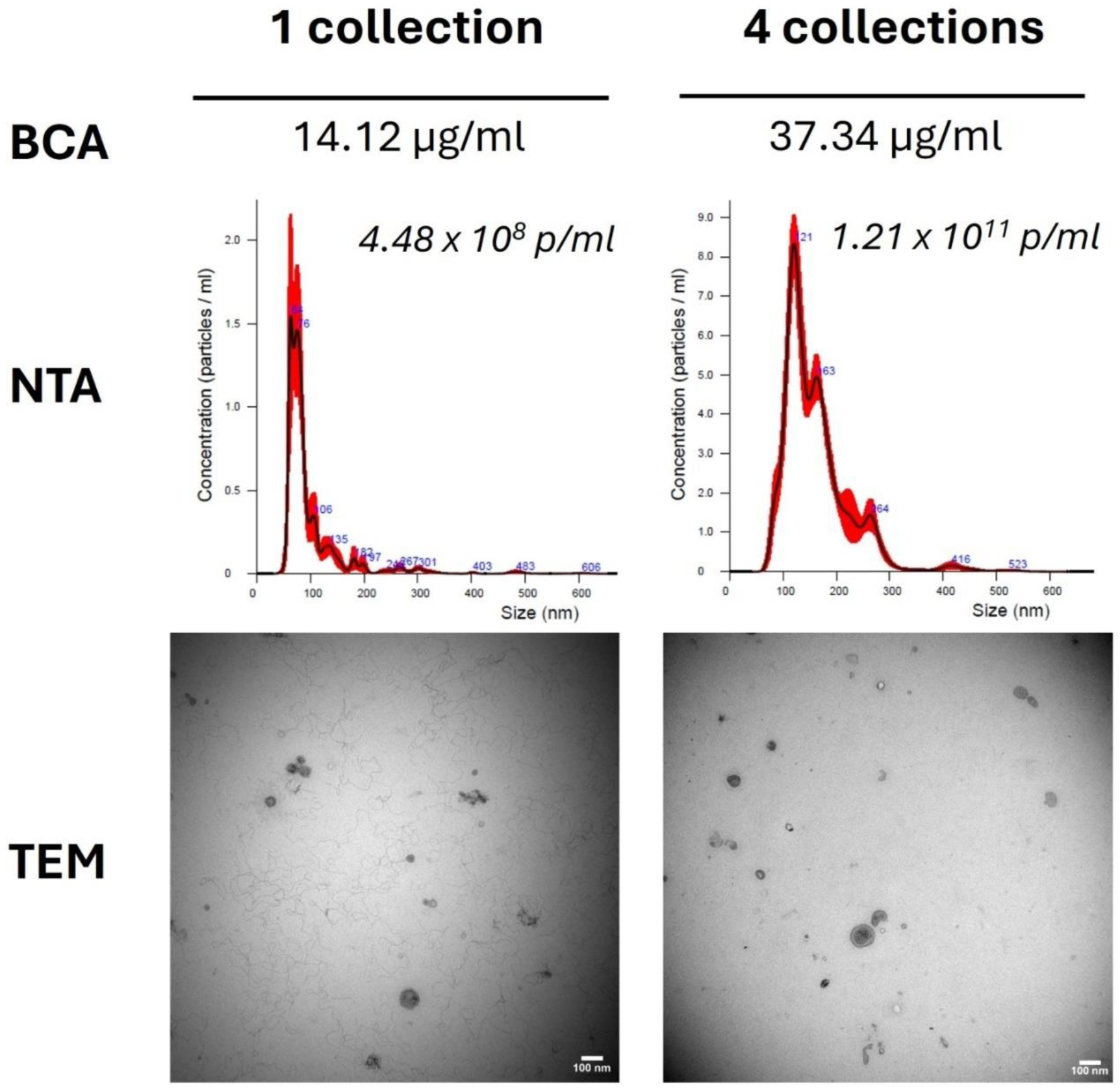
The effect of frequent *Leptolyngbya* medium collection on Lepto-EV yield. EV protein content, nanoparticle concentration, and TEM.

